# CpxR and HicB exert independent regulatory action on the gonococcal *hicAB*-encoded toxin-antitoxin system

**DOI:** 10.64898/2026.08.28.747762

**Authors:** Concerta L. Holley, Vijaya Dhulipala, William M. Shafer

## Abstract

The continued emergence of *Neisseria gonorrhoeae* (Ng) isolates resistant to front-line antibiotics has focused efforts on understanding how alternative therapies, such as the expanded use of gentamicin (Gen), might counteract this global public health problem. Focusing on Gen as a viable alternative antibiotic for the treatment of gonorrheal infections, we previously used RNA-seq to determine if sub-lethal levels of Gen might impact gonococci on a transcriptional level and showed that expression of the putative HicA-HicB toxin-antitoxin (TA) system was increased in response to sub-lethal Gen. Importantly, loss of this TA system resulted in reduction of Ng biofilm formation in a strain specific manner. Focusing on this strain specificity, we found that the CpxR/CpxA two-component system (TCS) influences expression of the *hicAB* operon independently of HicB autoregulation. We now report that CpxR selectively binds to the *hicAB* operon to enhance expression of *hicAB* but does not interfere with binding of HicB to the promoter region. Furthermore, we show that single base pair differences in the intergenic region between *hicA* and *hicB* impact regulation by CpxR. Hence, the regulation of the HicAB TA in gonococcal strains is a highly coordinated response that can involve autoregulation by HicB and the CpxRA TCS. We propose that this dual regulatory scheme maximizes the ability of Ng to respond to Gen and hostile environmental conditions.

**Author Summary:** Antibiotic concentrations within the body change over time as the drug is absorbed and spread through tissues and later eliminated. Therefore, during antibiotic treatment of disease, bacteria that survive the initial high dose exposure could be exposed to sub-lethal concentrations until the antibiotic clears. Currently, little is understood about the response of Ng to sub-lethal antibiotic concentrations; we believe closing this gap is crucial to combat antibiotic resistance and disease spread. We hypothesize that during antibiotic treatment, these sub-lethal concentrations could serve as a stress signal, allowing for internal changes that increase bacterial survival. In our previous work, we showed that sub-lethal levels of the aminoglycoside gentamicin can influence levels of the Ng HicAB toxin-antitoxin system. Here, we show that a two-component system CpxRA, a major stress response system in bacteria, directly influences expression of *hicAB*. Furthermore, we find that CpxR and HicB work synergistically, but independently, to regulate expression of the *hicAB* locus. Our study highlights the importance of understanding the complex interplay between regulatory systems and provides new insight into how Ng can use multiple stress responses simultaneously to survive the threat posed by antibiotics.

## Introduction

*Neisseria gonorrhoeae* (Ng) is a strict human pathogen that causes the sexually transmitted infection (STI) termed gonorrhea, which is the second most reported bacterial STI in the United States (U.S.) [1]. Ng causes localized, uncomplicated infections at genital mucosal surfaces as well as extragenital infections (e.g., oropharyngeal, rectal and eye). More invasive forms of disease in both females and males can occur and often result in severe consequences for the general and reproductive health of those infected individuals. Worldwide, it is estimated that 82 million cases of gonorrhea occur annually [2]. Unfortunately, Ng strains are becoming less susceptible or even resistant to ceftriaxone (CRO), which is the sole antibiotic now used in the U.S. and other countries for empiric, monotherapy of gonorrhea [3–5].

As a vaccine is not yet available to prevent gonorrhea, physicians rely on the efficacy of antibiotics to treat infections and reduce the spread of Ng in the community, making the prospect of untreatable cases of gonorrhea due to antibiotic resistance worrisome. To combat this troubling scenario, alternative drugs such as gentamicin (Gen) have been used in treatment regimens and clinical trials [6, 7]. Gen has been employed in East Africa (Malawi) for over 25 years and has been proposed for expanded use in treating gonorrhea in the U.S. [6, 8]. However, from a historical perspective, it is prudent to anticipate that the efficacy of Gen will be threatened by resistance development and global spread of resistant strains. We hypothesized that expanded use of Gen may promote the emergence of Ng with clinically relevant resistance and conducted earlier studies to understand how decreased susceptibility of Ng to Gen might develop [9, 10]. We found that Ng can develop a single amino acid change in *fusA* which encodes the bacterial translation factor elongation factor G, resulting in low-level Gen resistance [9, 10]. In addition to this mechanism of Gen resistance, earlier work showed that loss of the CpxRA, (also termed MisRS) two component system (TCS), rendered Ng hypersusceptible to aminoglycosides and cationic antimicrobial peptides. In addition, evidence has been presented that this TCS is important for regulation of Ng genes involved in iron acquisition [11], surface expression of vaccine candidates [12] and other genes involved in a general stress response [13, 14]. Interestingly, transcriptional profiling studies using different Ng strains (FA19 and FA1090) showed that the CpxR response regulator controlled expression of a lysogenic bacteriophage associated locus encoding a Type II toxin-antitoxin system (HicA-HicB), but only in strain FA1090. Moreover, incubation of Ng in the presence of sub-lethal levels of Gen enhanced expression of *hicAB* [15]. Importantly, deletion of *hicAB* in three Ng strains (FA1090, F62 and WHO-X), but not FA19, decreased biofilm forming capability [15]. Taken together, we hypothesized that *hicAB* expression in Ng involves two distinct regulatory schemes that can influence bacterial responses to antibiotics, environmental stresses, and elaboration of a virulence factor (e.g., biofilm formation).

## Results

### Strain-specific differences in the Ng *hicAB*promoter sequence influence biofilm formation and promoter activity

The Ng *hicAB* locus is within the genome of lysogenic prophage φ3 and is highly conserved across strains with potential binding sites for HicB and CpxR (Fig. 1). However, minor sequence differences exist in the intragenic region between *hicA* and *hicB* in strains FA19, F62 and FA1090 (Fig. 1). Briefly, compared to FA19, F62 contains a point mutation in the putative -35 region of the P2 promoter site (G to A) while FA1090 contains this G to A mutation and an additional mutation that inserts a C 16 bp upstream of the -35 region (Fig. 1). To understand if these intragenic sequence differences influence Ng biofilm forming activity seen previously in *hicAB* deletion mutants of Ng strains [15], we prepared complementing pGCC3 constructs that contained the entire *hicAB* locus from strains FA19 or F62. These plasmids were introduced into the *hicAB* deletion mutant of strain F62. As is shown in Fig. 2, deletion of *hicAB* in strain F62 containing pGCC3 alone had a severe biofilm defect compared to the parental strain. The defect was partially repaired by complementation using the *hicAB* sequence from F62 but not from FA19.

**Fig. 1.**
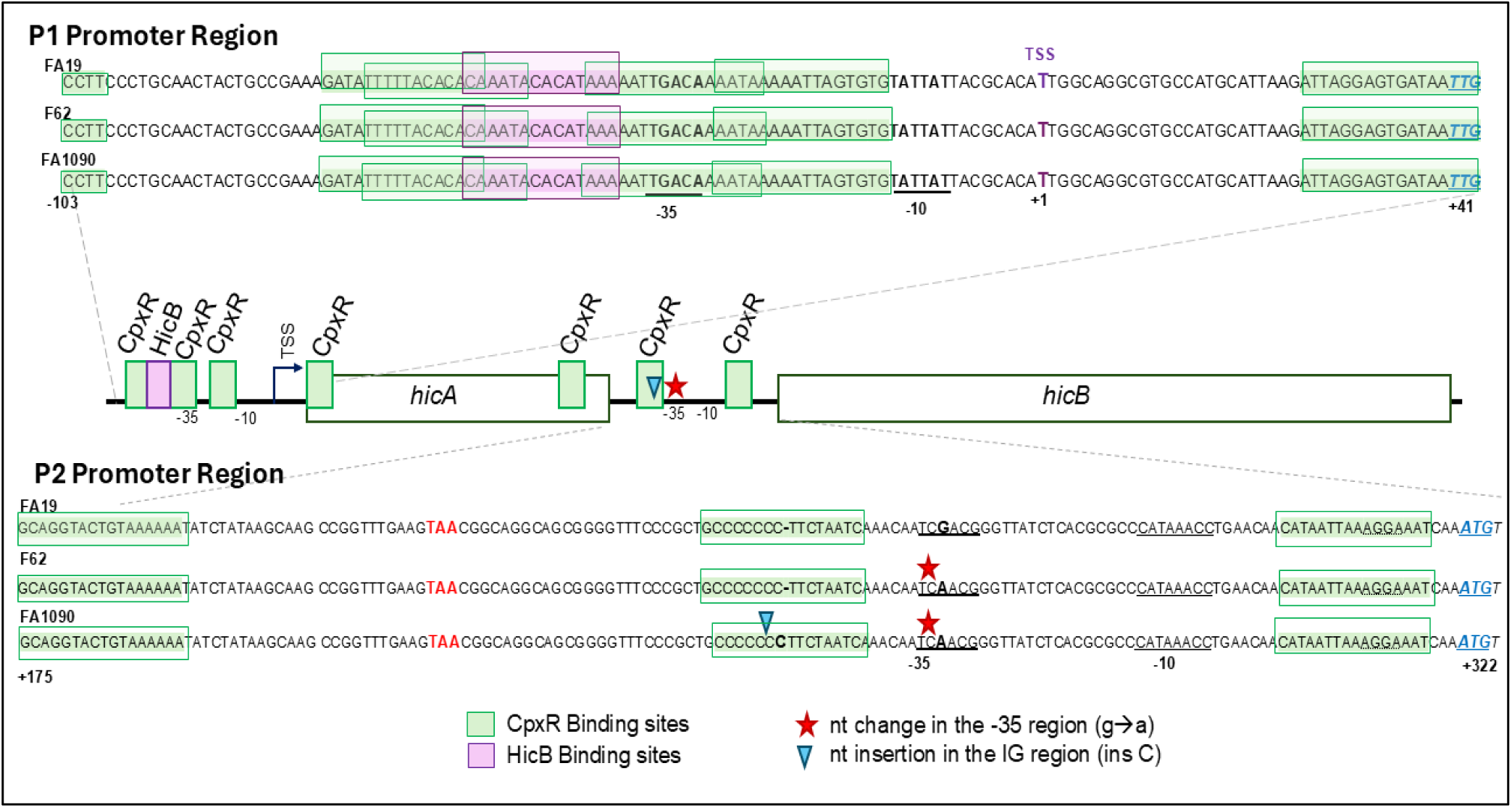
Sequence of the Ng *hicAB* operon. Shown is a comparison of the *hicAB* operon in Ng strains FA19, F62 and FA1090 with P1 (operon promoter, -103 to +41 relative to transcription start site (TSS)) and *hicA-hicB* P2 intergenic region (+175 to +322) expanded. SNPs are shown with an inverted triangle and an asterisk. HicB (purple) and CpxR(green) binding sites identified by foot printing are shown as filled rectangles (middle) and expanded sequence (open rectangle). The translational stop codon for *hicA* is shown as bold red text while the *hicA* and *hicB* translational starts are in underlined blue text. Final sequence matches were determined by location of protected region, sequencing ladder, and CpxR consensus sequence matching [13].

**Fig. 2.**
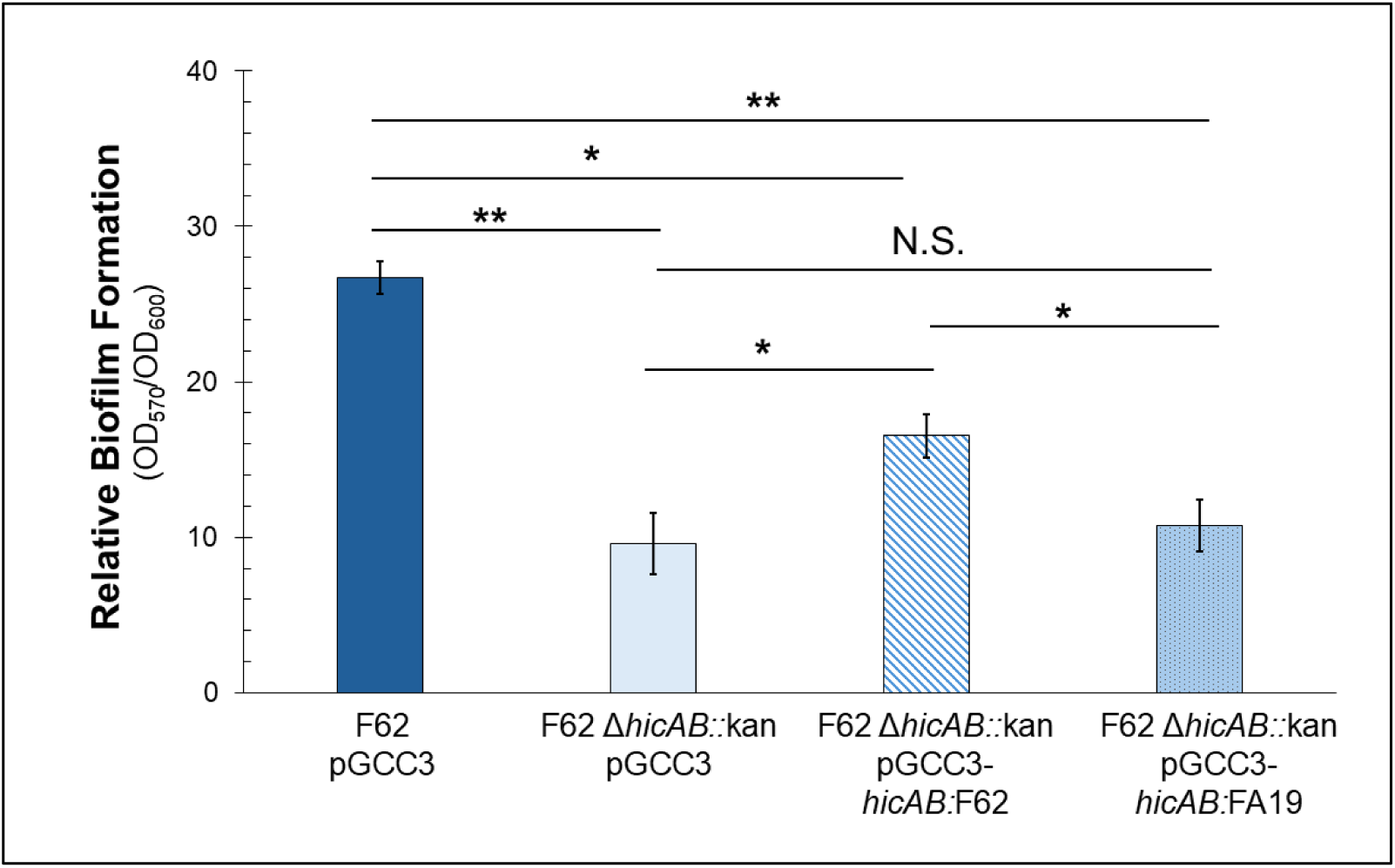
Minor differences in sequence impact biofilm formation and gene expression. Biofilm formation in strain F62 complemented with the *hicAB* locus from F62 and FA19 under the native promoter expressed frompGCC3. Data is shown as a ratio of crystal violet stained biomass (OD570) corrected by bacterial growth within each well (OD600) and are representative of at least four independent experiments. Error bars show standard error. *, *P ≤* 0.05; **, *P ≤* 0.01; N.S., not significant.

Based on the results from the cross-complementation study, we divided the full-length *hicAB* operon into its two potential promoter (P) regions. P1 contains the HicB binding site, transcriptional start site (TSS), and results in a *hicAB* transcript. P2 contains the intergenic region between *hicA* and *hicB* where the only sequence differences between strains FA19, F62, and FA1090 occur (Fig. 1). Using translational fusion strains grown in the presence or absence of 0.2X GEN MIC, we examined the P1 and P2 activity separately in the FA1090 WT, *hicAB* null mutant and complemented strains (Fig. 3A). We found that in the *hicAB* null mutant, there was significantly reduced P1 activity compared with the WT and complemented strains (Fig. 3B). Consistent with our earlier study [15], the addition of sub-lethal Gen increased promoter activity overall in the WT and null mutant strains. Of note, when using a translational fusion that lacks the HicB binding site (P1ΔS1), and thus removing HicB repression of the promoter, significantly more activity at P1 compared to the WT was observed in the presence of Gen (Fig. 3B). Moreover, there was significantly less activity at P2 in the *hicAB* null mutant compared to the WT strain in the absence of Gen (Fig. 2C). Interestingly, complementation led to significantly increased activity at P2 but only in the absence of Gen. There was no difference in P2 activity between the WT, null mutant, or complement strain in the presence of Gen (Fig. 3C). However, when P2 from FA19 was expressed in the FA1090 strain background, a response to Gen was observed (Fig. 3C). This led us to directly compare activity of the strain-specific promoter regions. We found that promoter activity from strains containing the FA19 sequence had reduced activity compared to the FA1090 sequence for both P1 and P2 (Fig. 3D). Therefore, P1 is likely the primary Gen responsive promoter in Ng and the primary location of *hicAB* autorepression by HicB.

**Fig. 3.**
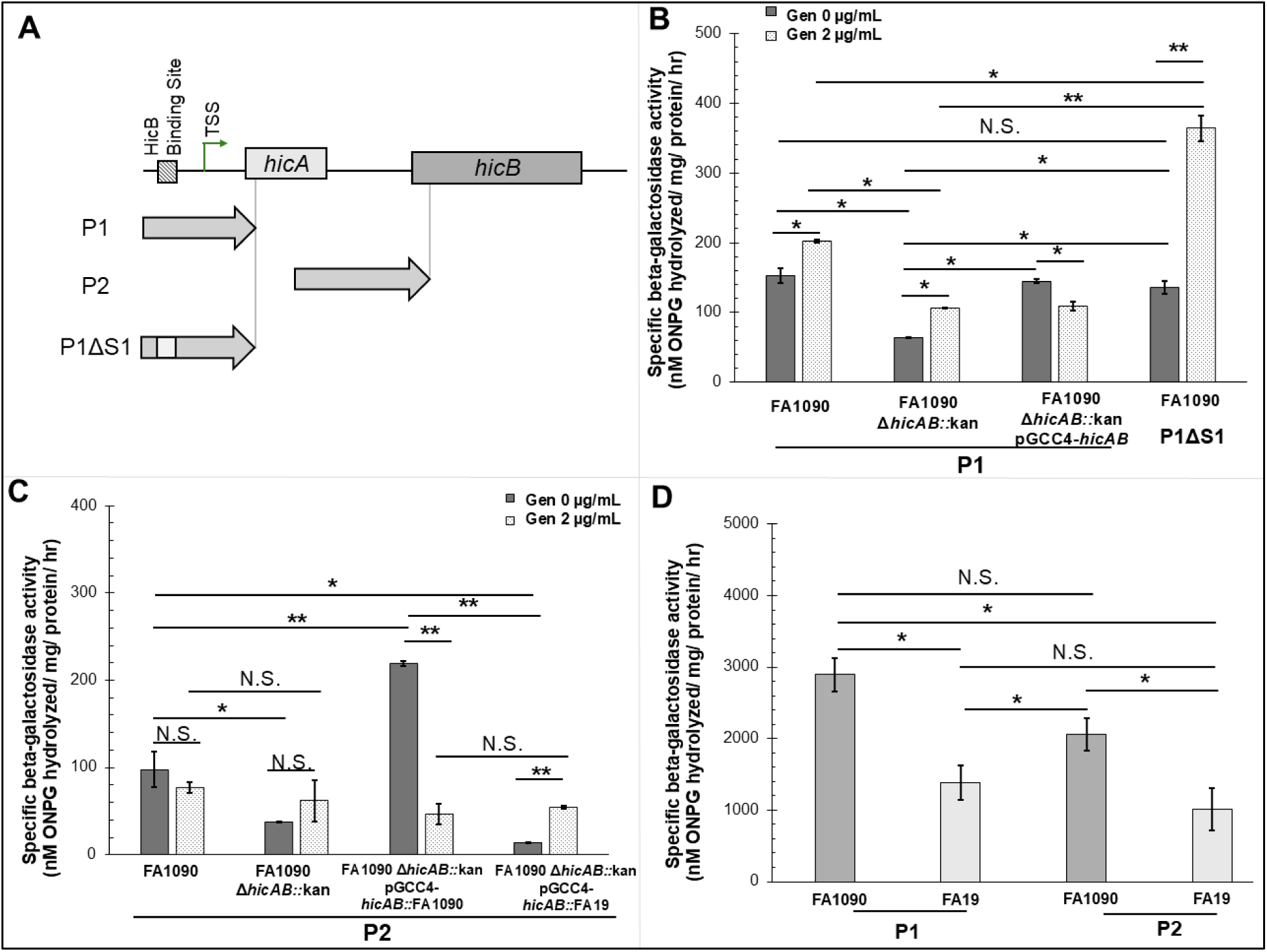
Sub-lethal Gen exposure differentially regulates the primary and secondary *hicAB* promoters. A) Schematic of the locations of P1 and P2 relative to *hicA* and *hicB.* The approximate location of the HicB-binding site and the Transcription Start Site (TSS) are indicated. Sizes are not to scale. B) The specific β-galactosidase activity per mg of total protein in cell extracts of reporter strains containing the P1*-*lacZ or P1ΔS1-lacZ fusions in the FA1090, *hicAB* null mutant and complement strains in the presence or absence of Gen 2 µg/ml C) β-galactosidase activity of reporter strains containing the FA1090::P2*-*lacZ fusions in the FA1090, *hicAB* null mutant and complement strains or FA19::P2-lacZ in FA1090 in the presence or absence of Gen 2 µg/mL. D) β-galactosidase activity of reporter strains containing the FA1090/FA19 P1-lacZ or FA1090/FA19 P2-lacZ fusions in FA1090. Data are representative of at least four independent experiments. Error bars show standard error. *, *P ≤* 0.05; **, *P ≤* 0.01; N.S., not significant.

### HicB regulates *hicAB* through binding to P1 region

We assessed if HicB autoregulation of *hicAB* requires both P1 and P2. Consistent with our previous work [15], using an electrophoretic mobility shift assay (EMSA) we found that HicB binds P1in a specific manner as competition with an unlabeled sequence lacking the HicB binding site or a non-specific probe did not disrupt HicB binding to P1 (Fig. S1A). In contrast, under the conditions employed we did not detect binding HicB to P2(Fig. S1B). As the P2 is not Gen responsive and HicB does not interact with it, we hypothesized that regulation of P2 must occur primarily through an alternate independent mechanism.

### CpxRA regulation of *hicAB*

We next sought to determine what transcriptional factors other than HicB contribute to control *hicAB* expression. We hypothesized that the CpxRA TCS could influence *hicAB* expression, possibly in a strain-specific manner given that RNA-seq analysis of CpxR mutants in strains FA19 and FA1090 revealed overlapping yet distinct gene sets [13, 14]. Of note, in FA1090, but not FA19, loss of *cpxR* resulted in decreased expression of the *hicAB* operon. We first confirmed the RNA-seq data and showed that *hicA* and *hicB* levels were significantly reduced in the FA1090 *cpxR* mutant but not in the FA19 mutant (Fig. 4A). Furthermore, in the *hicAB* null mutant, *cpxR* transcripts were significantly increased compared to the WT strain, which likely is the reason why the level of the CpxR-activated *ompA* gene transcript was increased in the mutant [12]. Critically, complementation of the mutant with *hicAB* returned *ompA* and *cpxR* transcript levels to WT levels (Fig. 4B). We next examined biofilm formation in the CpxR null mutants. Loss of *cpxR* reduced biofilm formation in the FA1090 background but not in the FA19 background (Fig. S2); the reduction in biofilm formation could be restored by complementation with WT *cpxR.* Given that loss of *hicAB* also impacts biofilm formation in a strain-specific manner and that there are differences in promoter activity linked to differences in sequence, we hypothesized that CpxR is involved in regulation of *hicAB*.

**Fig. 4.**
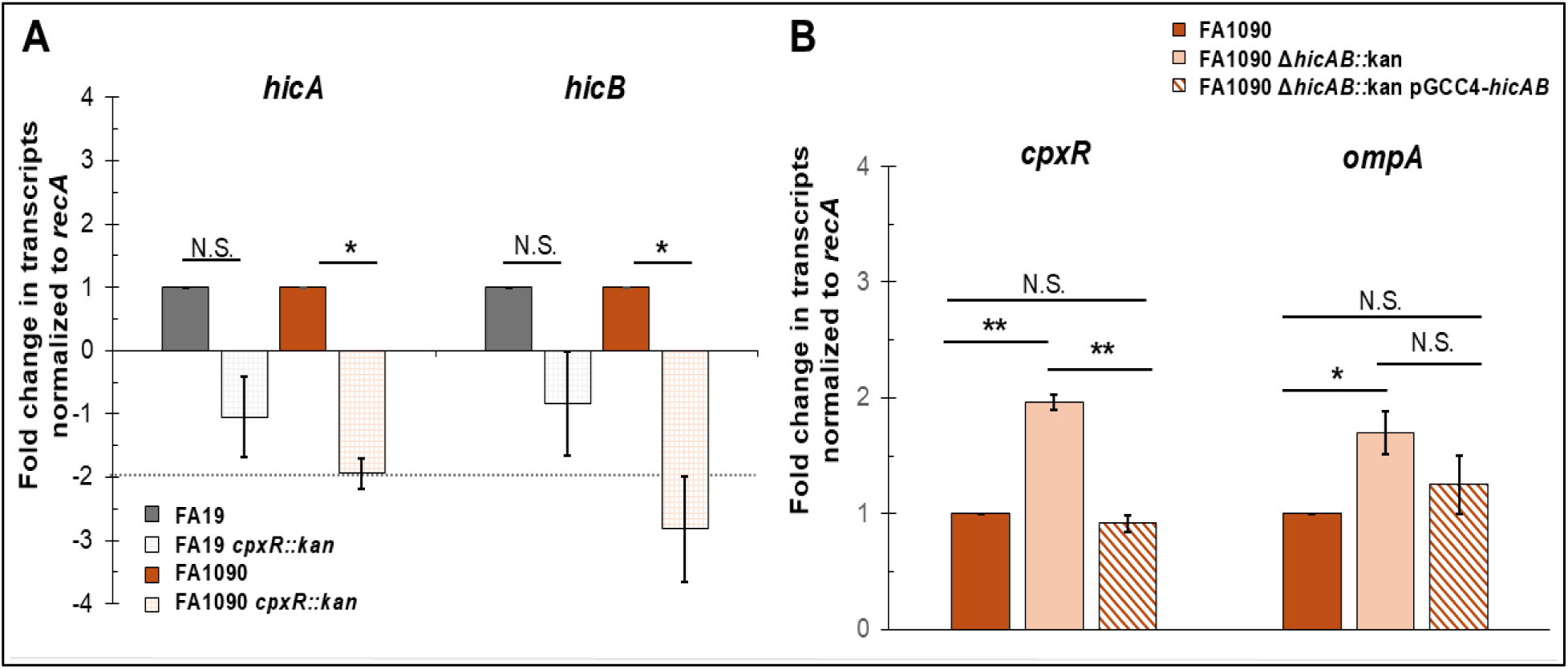
Loss of *cpxR* impacts *hicAB* expression in FA1090 but not FA19. A) Expression of *hicA* and *hicB* transcripts FA19, FA1090 and their isogenic *cpxR* null mutants. The dotted line indicates the corresponding RNA-seq cutoff values for differentially expressed genes. qRT-PCR analysis shows reduced transcripts of both *hicA* and *hicB* in the FA1090 background only. B) Loss of *hicAB* increases expression of *cpxR* in FA1090. Transcript levels of CpxR regulated gene *ompA* were increased in the *hicAB* null mutant. Complementation restored transcripts levels to WT. Data are representative of at three independent experiments. Error bars show standard error. *, *P ≤* 0.05; **, *P ≤* 0.01; N.S., not significant.

### CpxR directly regulates expression of *hicAB*

Upregulation of *hicAB* expression was linked to exposure to sub-lethal Gen [15] and CpxR null mutants show increased susceptibility to aminoglycosides including Gen [13]. For this reason, we next determined if the presence of sub-lethal Gen would impact expression of the *hicAB* locus in *cpxR* null mutants. We first examined the impact of loss of *cpxR* on transcription of *hicA, hicB* or the *hicAB* bicistronic operon in increasing concentrations of Gen. An RT-PCR was performed using RNA isolated from FA1090 or FA1090 *cpxR*::kan grown on agar containing 0.25 and 0.5 X the Gen MIC (MICs are 8 and 1.25 µg/mL respectively). In FA1090, *hicB* and the *hicAB* operon transcripts were reduced in 0.5X Gen MIC compared to 0 or 0.25X (Fig. S3, left panels). In the *cpxR*::kan mutant, there was no such reduction in transcripts with increasing Gen (Fig. S3, right panels). Taken together, this suggests that the WT and *cpxR::*kan strains have different transcriptional responses to sub-lethal Gen.

Given this result, we next examined promoter activity in the WT and *cpxR::*kan mutants. We found that not only did loss of CpxR significantly increase P1 activity, but that activity was also increased in the presence of sub-lethal Gen for both strains (Fig. 5A). In contrast, P2 activity was reduced in the presence of sub-lethal Gen and in the absence of CpxR (Fig. 5B). Thus, CpxR represses P1 activity while enhancing P2 activity. Deletion of both *hicAB* and *cpxR* significantly reduced activity of both promoters consistent with the idea that both systems are needed for optimal P activity. To further understand the role of CpxR regulation of *hicAB*, we determined if phosphorylation of CpxR was necessary for promoter activity. Phosphorylation at a conserved aspartic acid residue (D51 in *E. coli,* D52 in Ng) is crucial for CpxR function [16]. We analyzed promoter activity in strains containing either the WT or D52A CpxR complemented strains and found that phosphorylation was necessary for activity of both P1 (Fig. 5C) and P2 (Fig. 5D). Therefore, phosphorylated CpxR is likely necessary for repression of P1 and activation of P2.

**Fig. 5.**
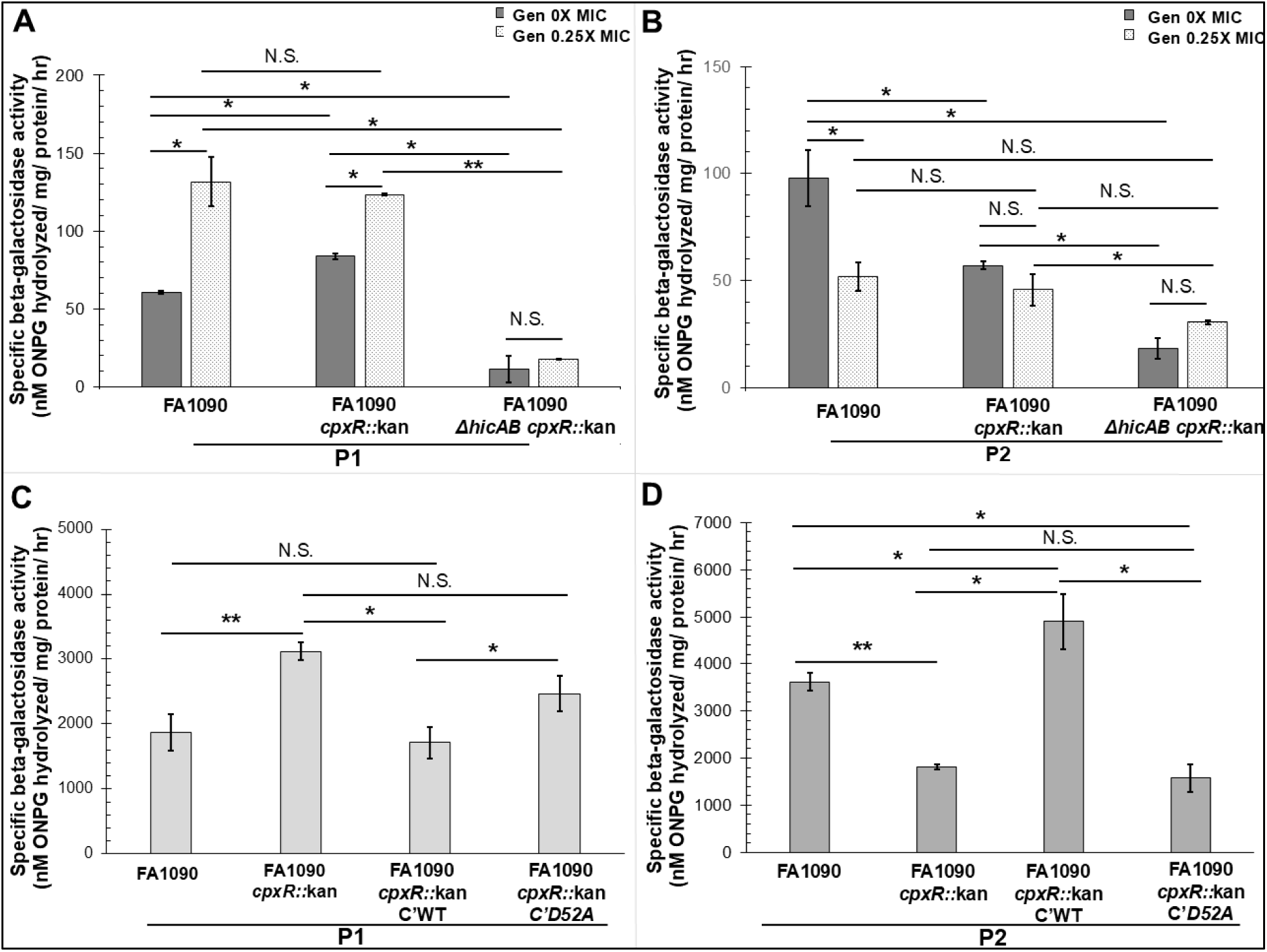
Sub-lethal Gen exposure differentially regulates *hicAB* P1 and P2 . A and B) The specific β-galactosidase activity per mg of total protein in cell extracts of reporter strains containing the P1*-lacZ* (A) or P2-*lacZ* (B) fusions in the FA1090, *cpxR* null and *cpxR hicAB* double null strains in the presence or absence of Gen 0.25X Gen MIC (2 and 0.03125 µg/mL respectively). C and D) β-galactosidase activity of reporter strains containing the P1*-lacZ* (C) or P2-*lacZ* (D) fusions in the FA1090, *cpxR* null mutant and either WT (C’WT) or phosphorylation null (C’D52A) complement strains. Reporter constructs are located at an alternate site within the genome. Data are representative of at least four independent experiments. Error bars show standard error. *, *P ≤* 0.05; **, *P ≤* 0.01; N.S., not significant.

To identify the sequence of the target DNA capable of binding CpxR, we used a DNase I protection assay. As shown in Fig. 1, predicted CpxR binding sites are located at multiple locations in or adjacent to both P1 and P2. These sites include three sites in the *hicA-hicB* intergenic region (P2), five sites upstream of the TSS (P1) and two sites within the coding region of *hicA* (Fig. 1). Detailed DNase I foot printing figures are shown in Fig. 6 (P2) and Fig. S4 (P1). The results showed overlap of the HicB binding site and two of the CpxR binding sites in P1 which was visible in the DNase I protection analysis (Fig. S4). For P2, one of the CpxR binding sites is located upstream of the putative -35 region, overlapping the region where the sequence differs between FA1090 and FA19/F62 (Fig. 6). The insertion of the C nucleotide reduces the predicted matches to the published CpxR consensus sequence [13] from 73% in FA19/F62 to 66% in FA1090; as a result, the CpxR binding site shifts slightly in the FA1090 sequence. Taken together, we propose that the Ng CpxR binds the *hicAB* promoter in regions that both overlap and are distinct from that of HicB.

**Fig. 6.**
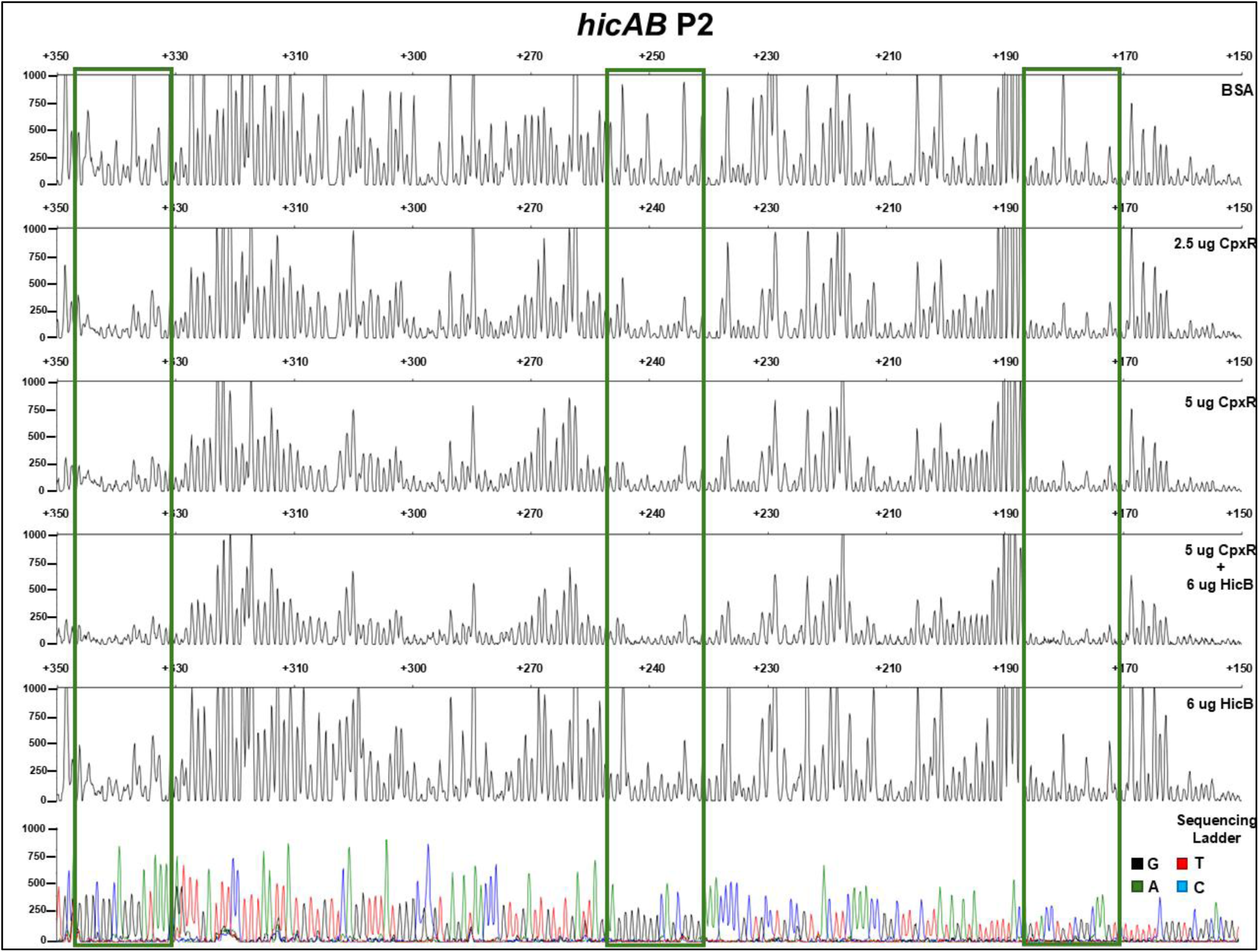
CpxR protects multiple regions of *hicAB* P2. A DNA fragment spanning *hicAB* P2 region from nucleotide +350 to +150 (relative to the TSS) was fluorescently labeled with 6-FAM (coding strand) and HEX (template strand) and incubated with BSA (control reaction), CpxR or HicB prior to digestion with DNase I. The DNase I digestion products were analyzed by capillary electrophoresis. The fluorescence signal corresponding to the HEX probe is shown on the y axis of each electropherogram. Fragment coordinates (relative to the TSS) are shown on top. Electropherograms corresponding to increasing concentrations of BSA control, CpxR and/or HicB are shown. The CpxR protected regions are boxed in green. Sequencing reactions were manually generated (bottom panel). DNA sequence of the CpxR-protected region is shown in Fig. 1.

## Discussion

Interactions between two component (TCS) and toxin-antitoxin (TA) systems are often directly linked to virulence or persistence phenotypes. When a TCS senses environmental stress, such as the presence of antibiotics, it can trigger alteration of specific TA modules to fine-tune survival efforts. CpxRA TCS has been linked to several TA systems in response to environmental triggers. For example, deletion of the *S*. *enteritidis* CpxR induced multiple TA systems in response to the polycationic antimicrobial agent, chlorhexidine [17]. The *E. piscicida* HigA antitoxin was upregulated by CpxR activation in the presence of serotonin, a condition that ultimately leads to disturbance of the host immune response [18]. In *B. subtilis*, the DegS/DegU TCS regulates multiple TA operons involved in virulence and biofilm formation [19]. CpxR directly controls *hha* promoter activity in *P. mirabilis* to induce *hha* protein toxin expression which controls virulence and biofilm formation [20]. Although Hha is not part of a TA system, activity of CpxR on its promoter directly impacts virulence and biofilm formation, highlighting the importance of CpxRA in bacterial pathogenesis.

With the results presented within this work and our previous study [15], we propose a model (Fig. 7) to describe the interaction between CpxRA and HicAB in response to sub-lethal Gen exposure. We hypothesize that sub-lethal Gen induces mistranslation leading to an accumulation of misfolded proteins in the periplasmic space. CpxA senses this build-up causing it to phosphorylate CpxR triggering transcriptional regulation of target genes such as the *hicAB* operon. Under normal conditions, the HicA toxin is neutralized by HicB antitoxin. HicB also represses transcription through autoregulation of the *hicAB* operon promoter. When activated, HicB is degraded releasing HicA to cleave target mRNAs. Thus, increasing production of the HicAB TA system by CpxRA would prime the cell for initiation of a ‘persister’ state that can rapidly respond to environmental triggers. Taken together, CpxRA and HicAB work synergistically for gonococcal persistence amidst antibiotic pressure.

**Fig. 7.**
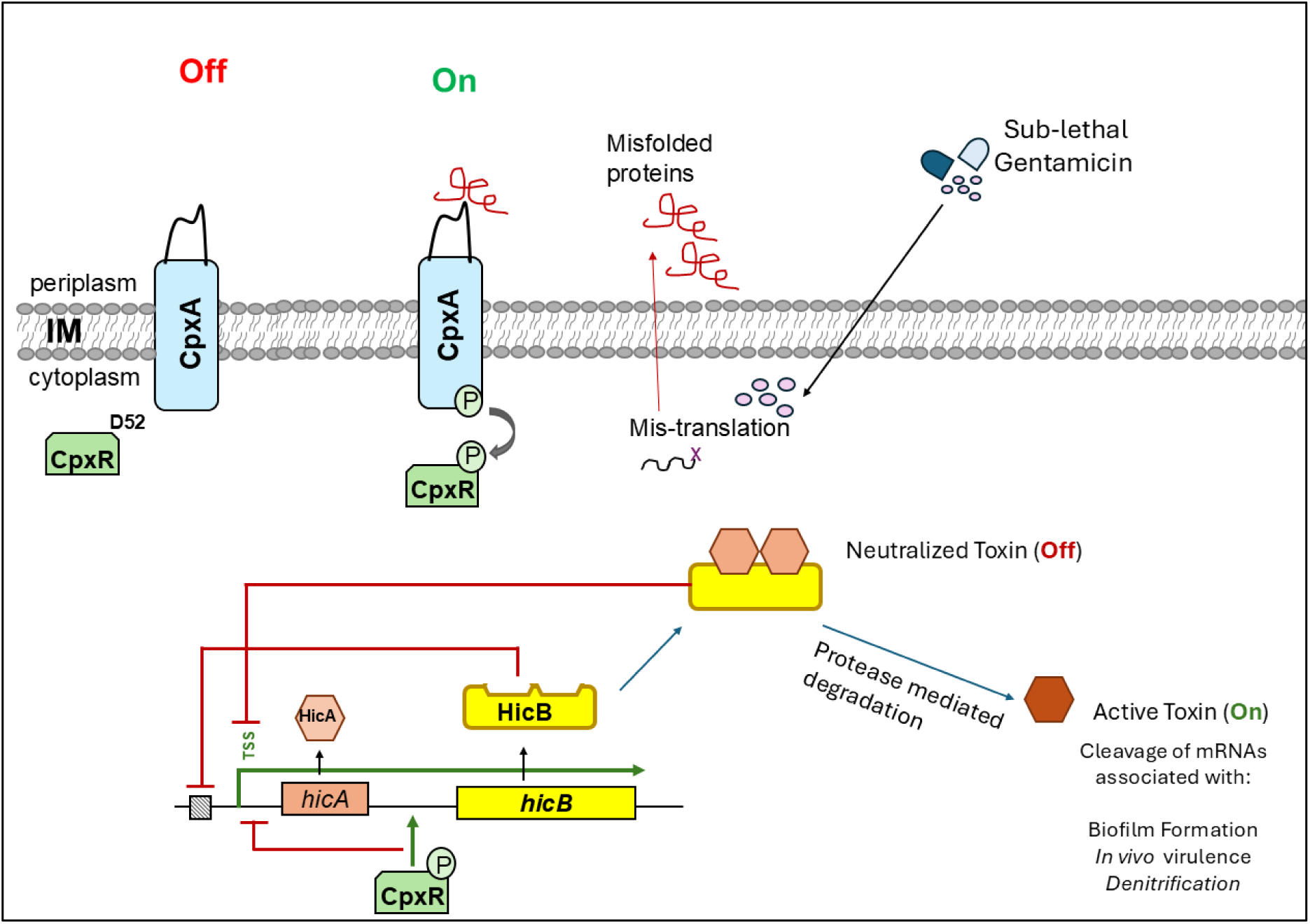
Model of HicA-HicB Toxin-Antitoxin regulation in Ng. Sub-lethal Gen may trigger mistranslation which leads to mis-folded protein accumulation. Sub-lethal Gen upregulates expression of the HicAB TA system in response to the environmental stress. Activation of the HicAB TA system is related to several survival processes including biofilm formation, *in vivo* virulence, and denitrification. In steady conditions, HicB binds to its own promoter to function as a repressor and inhibit transcription of *hicA*. In stress conditions, HicB is degraded and free HicA toxin is free to degrade target mRNAs while HicB is no longer capable of repressing transcription. Likewise, in normal conditions, the Ng CpxRA TCS system is “Off”. CpxA senses accumulation of misfolded proteins in the periplasmic space and activates CpxR by phosphorylation (On). CpxR can inhibit transcription from P1 by inhibiting activity of the TSS potentially by blocking transcription of *hicA*. CpxR can also increase activity of P2 by binding to the region and potentially recruiting RNAP to preferentially transcribe *hicB*.

The *in vivo* trigger(s) for the gonococcal CpxRA are l unclear [13, 14]. However, our earlier work showed that loss of CpxRA resulted in hyper-susceptibility of Ng to cationic antimicrobials such as aminoglycosides (including Gen) and antimicrobial peptides [13]. CpxR has been shown to be important for resistance to antibiotics [21]. Additionally, moderate induction of CpxR was associated with increased drug tolerance by strengthening the cell wall [22]. We propose that antimicrobial exposure could be a natural *in vivo* trigger for the gonococcal CpxRA system; careful study of the CpxRA response to host-derived antimicrobials and clinically used antibiotics could further elucidate the role of CpxRA in Ng pathogenesis.

This is not the first report of the gonococcal CpxRA differentially regulating genes depending on environmental conditions. CpxR differentially regulates transferrin binding proteins *tbpB* and *tbpA* depending on the environmental availability of iron [11]. In iron-replete conditions, CpxR represses transcription of the operon working cooperatively with Fur protein; under depleted condition, cpxR binds to multiple locations in the *tbpBA* promoter to maximize expression. We suggest that CpxR functions similarly with the *hicAB* operon, working with HicB to repress operon (e.g., *hicA*) transcription in neutral conditions while increasing *hicB* single gene transcripts and de-repressing *hicA* in stress (antibiotic) conditions. We note that the spacing between the proposed -35 and -10 of P2 is a suboptimal 16 bp [11] and similarly to the *tbpBA* operon, CpxR may compensate for the suboptimal promoter elements and assist interaction between *hicAB* transcript and RNAP.

In *E.coli*, the *hicAB* operon contains two promoters (ECP1 and ECP2) that generate two independent TSS. ECP1 allows expression of both the toxin and antitoxin genes while ECP2 only allows transcription of antitoxin [23]. In this way, *E. coli* can maintain a balanced toxin-antitoxin ratio that can respond to the changing environment rapidly. However, while a dual TSS *hicAB* promoter has only been shown in *E. coli* [24] we have no evidence for a second TSS in Ng using an *in vitro* transcription assay. Further, although there are strain-specific phenotypes, the only sequence differences we could detect in *hicAB* locus are in P2 region. CpxR preferentially binds to P2 but can interact with elements in both P1 and P2. The binding of HicB to P1 may allow CpxR to interact with P2 more strongly; conversely, CpxR interaction with the promoter may cause HicB to bind to a heretofore unidentified second site. Our collective DNA binding results suggest that CpxR and HicB binding to target DNA sequences is not mutually exclusive or antagonistic; in fact, the presence of both CpxR and HicB is associated with more robust binding and increased promoter activity. We propose that HicB and CpxR work cooperatively in regards to *hicAB* transcription at least in the FA1090 strain background.

In conclusion, we posit that transcriptional regulation of the *hicAB* TA system mediated by CpxRA is an important part of the gonococcal antibiotic stress response. We propose that the study of gonococcal responses to sub-lethal antibiotic exposure can provide insights regarding transcriptomic and physiologic changes bacteria undertake during natural infection and treatment.

## Materials and Methods

### Bacterial strains, plasmids, and primers

Ng strains and their isogenic genetic derivative strains, along with the plasmids used and their *Escherichia coli* hosts, are listed in Supplementary Table S1. The oligonucleotide primers used in this study are listed in Supplementary Table S2. *E. coli* strains were routinely cultured on Luria-Bertani (LB) agar or in LB broth (Difco, Sparks, MD) containing 50 µg/ml Kanamycin, 100 μg/ml ampicillin, or 100 μg/ml chloramphenicol, as necessary. Gonococci were grown on gonococcal base (GCB) agar (Difco, Sparks, MD) containing Kellogg’s supplements I and II at 37°C under 5.0% (v/v) CO2 [25] containing 50 µg/ml Kan, 1 μg/ml erythromycin, or 1 μg/ml chloramphenicol as necessary. Liquid cultures of gonococci for growth assays were begun by inoculating plate-grown cells in pre-warmed GCB broth containing Kellogg’s supplements I and II and 0.043% (w/v) sodium bicarbonate and grown in a 37°C water bath with shaking. Gentamicin was added to concentrations specified in the text and appropriate for each strain and condition being evaluated.

### Complementation of the Δ*hicAB::kan* mutants

Strains were complemented as previously described using the pGCC3 or pCGG4 complementation vector [26]. Construction of the pGCC4-*hicAB* plasmid is described elsewhere [15]. For construction of pGCC3-*hicAB,* the entire *hicAB* operon and flanking regions containing the promoter regions were amplified from FA19 or F62 genomic DNA using primers GCC3 hicABF2 and GCC3 hicABR2 inserted into digested pGCC3 vector. The resulting plasmid was used to transform Δ*hicAB::kan* mutants. Colonies were selected on chloramphenicol 1 μg/mL and transformants were verified by PCR and sequencing using primers lctp and ermCRev.

### Analysis of Ng transcripts

For measurement of target gene expression, gonococci were harvested at late-log phase, and the pellets were stored at −70°C. RNA was purified by Trizol extraction as per manufacturer instructions (Thermo Fisher Scientific, Waltham, MA) followed by Turbo DNA-free (Ambion, Austin, TX) treatment. cDNA was generated using a QuantiTect reverse transcriptase kit (Qiagen, Venlo, Netherlands). We validated our qRT-PCR methods by examining primer efficiency, primer specificity (melt temperature), and linear dynamic range for each primer pair utilized herein. For qRT-PCR analysis, the normalized expression of each target gene was calculated using *recA* as a housekeeping reference gene [27]. All qRT-PCRs were performed in technical and biological triplicates. For RT-PCR, 2 µg of RNA was collected from strains grown on GCB agar with 0, 0.25 and 0.5 X the respective strain Gen MIC. RNA was isolated as described above. cDNA was generated using the SuperScript™ IV First-Strand Synthesis System (Thermo Fisher Scientific, Waltham, MA). cDNA was then diluted 1:100 and used as template in a standard PCR reaction.

### Purification of recombinant protein

Purification of recombinant HicB and CpxR are described previously [15, 28]. Briefly, both CpxR and HicB were purified as per the manufacturer’s protocol using a (Ni+2 -NTA) column (Millipore Sigma, Burlington, MA). Protein was eluted in buffer containing 250 mM imidazole, dialyzed to remove imidazole using 10 mM PBS (137 mM NaCl, 2.7 mM KCl, 10 mM Na2HPO4, 2 mM KH2PO4), and concentrated. Dithiothreitol (DTT) and glycerol were added to a final concentration of 1 mM and 10 % (v/v), respectively. The purity of recombinant protein was confirmed by SDS-PAGE electrophoresis and staining with Coomassie blue.

### EMSA for detection of CpxR and HicB binding to target DNA

For the full-length probe, the *hicAB* operon including the promoter and intergenic region was amplified by PCR from FA1090 genomic DNA using the primers hicAB_P1_For and hicAB_P2_Rev. For the P1 probe, the promoter region in front of *hicA* was amplified by PCR from FA1090 genomic DNA using the primers hicAB_P1_For and hicAB_P1_Rev. Site-directed mutagenesis was used to delete the HicB binding site. The P2 probe was amplified using primers hicAB_P2_F2 and hicAB_P2_Rev. EMSAs were conducted using the second-generation digoxigenin (DIG) gel shift kit (Roche Applied Sciences, Madison, WI) as previously described [29]. P1, P2 and FL probes were labeled with DIG following manufacturer’s protocol. Specificity of CpxR binding was assessed by adding 100-fold excess of unlabeled DNA fragments encoding either the specific competitor (P1, P2, FL or P1ΔS1) or a non-specific DNA competitor (rnpB) with purified protein for 30 min at 30°C in Binding Buffer: 20 μL of 20 mM Hepes, pH 7.6, 1 mM EDTA, 10 mM (NH4)2SO4, 1 mM DTT, 0.2% (v/v) Tween-20, 30 mM KCl and 1.25 ng/μL Type XV calf thymus DNA. Reactions were separated by electrophoresis in 5% Mini-Protean TBE Precast Gels (Biorad, Hercules, California) and transferred to Zeta-Probe membranes (Biorad, Hercules, California) and crosslinked. Blots were developed using an anti-DIG Fab fragment-AP conjugate (Roche Applied Sciences, Madison, WI).

### DNase I Protection Assays

A PCR fragment spanning the *hicAB* locus from nucleotide-253 to +448 (relative to the Transcription Start Site) was amplified using the 6-carboxyfluorescein (FAM)- and 6-carboxy-2’,4,4’,5’,7,7’-hexachlorofluorescein (HEX)- labeled primers [FAM] hicAB_FP_For and [HEX] hicAB_FP_Rev. CpxR and HicB protein binding to the labeled DNA probe and DNase I digestion reactions were performed as described previously [30] using phosphorylated CpxR [13]. Detection of the DNase I digestion peaks was conducted in a 3730 capillary sequencer (Applied Biosystems, Waltham, MA) by Azenta Life Sciences (Chelmsford, MA) and the alignment of the corresponding electropherograms was generated using PeakScanner v. 2.0 (Applied Biosystems, Waltham, MA). Negative control reactions were done using BSA at the same max concentration used for CpxR. Dideoxy sequencing reactions were manually generated using the HEX primer and a PCR fragment encoding the *hicAB* operon. The final electropherograms of the sequencing reactions were horizontally aligned with those generated in the footprint.

### β-galactosidase (β-gal) assays

Several plasmids containing *hicAB* promoters translationally fused to the truncated, promoter-less *lacZ* gene in pLES94 were constructed. Primers hicAB_P1_For and hicAB_P1_Rev were used to amplify the P1 region and primers hicABP2_F2 and hicAB_P2_Rev were used to amplify the P2 region. The P1 region contains nucleotide -137 to +49 (relative to the Transcription Start Site) and the P2 region contains nucleotide +37 to +336. The plasmids, pLES94-P1-lacZ and pLES94-P2-lacZ were used to transform FA1090, FA1090 Δ*hicAB::kan* and FA1090 *cpxR::kan,* and FA1090 Δ*hicAB::kan cpxR::ermC.* Gonococcal transformants were selected on GCB agar containing 1 μg/mL of chloramphenicol and further verified by PCR and sanger sequencing. SDM was utilized to delete the previously identified HicB binding site [15] using primers P1S1delF and P1S1delR to create ples94-P1ΔS1-lacZ. Strains containing *lacZ* translational fusions were grown overnight on GCB agar plates containing 1 μg/mL of chloramphenicol. β-gal assays were conducted as previously described. Briefly, cells were scraped from GCB agar into 1 ml of PBS, pelleted, washed, and resuspended in cold 1X Z buffer (60 mM Na2HPO4 · 7H2O, 40 mM Na2HPO4 · H2O, 10 mM KCl, pH 7.0). Cells were lysed by three freeze-thaw cycles and pelleted at 10,00 x g for 10 mins. 230 ul of the lysate was added to a mixture containing 4 mg/ml *o*-nitrophenyl-β-galactopyranoside and 50 mM β-mercaptoethanol in Z buffer. The samples were incubated for 24 hours at 37°C and the reaction was terminated by the addition of 500 μl 1 M Na2CO3. β-galactosidase activities are given in Miller units using the formula [1,000 × OD420nm / (*t* × *v* × OD600nm)], where *t* is the reaction time in min and *v* is the volume of cell lysates in mL per reaction.

### Analysis of Ng biofilms

Briefly, 5×105 of each gonococcal strain was seeded into 96 well microtiter plates (Costar, Corning, New York) and incubated at 37°C for 24h to allow static biofilm formation. Bacterial density was then read (absorbance at 600 nm) using a Victorx3 2030 Multilabel Reader (Perkin Elmer, Waltham, MA) to determine the baseline growth for each well. After incubation, planktonic bacteria were removed, and the wells were gently washed 3 times with phosphate-buffered saline. Plates were allowed to dry and then stained for 30 mins with 0.1% (w/v) crystal violet (CV). Plates were washed with PBS, dried, and CV-stained biofilms were solubilized with DMSO. Absorbance, indicative of biofilm biomass, was read at 570 nm. Data were adjusted for background and each assay was performed using triplicate wells in at least three independent experiments. Relative biofilm formation was calculated as ratio of the OD570/ OD600 for each well and averaged across strains.

### Statistical Methods

All the data were expressed as means with standard deviation (SD). Statistical significance between all quantitative data was analyzed by Student t-tests or one-way ANOVA followed by Tukey’s honestly significant difference posthoc test. Statistical significance was set at P < 0.05.

## Data availability Statement

The datasets that supported the findings of this study are available from the corresponding author upon request. The supporting raw data for this manuscript is available in the Harvard Dataverse under https://doi.org/10.7910/DVN/NFDDTC.

## Acknowledgements

This work was supported by NIH grants R01 AI147609, R01 AI021150, and P01 AI197193 (W.M.S.). W.M.S. is the recipient of a Senior Research Career Scientist Award (IK6BX005390) from the Biomedical Laboratory Research and Development Service of the U.S. Department of Veterans Affairs. The contents of this article are solely the responsibility of the authors and do not necessarily reflect the official views of the National Institutes of Health or the U.S. Department of Veterans Affairs. The authors have no competing interests to declare. The authors have no conflicts to report.

